# An increased number of heterozygous calls in the Axiom™ Equine Genotyping Array

**DOI:** 10.1101/2025.09.16.676615

**Authors:** Annik Imogen Gmel, Ali Pirani, Liz McInnis, Markus Neuditschko

## Abstract

Single nucleotide polymorphism (SNP) arrays are commonly used in livestock genetics to investigate complex traits including genome-wide analysis and fine mapping, genomic prediction, genetic diversity and selection signature analyses. In the context of a European Equine diversity study, we analysed the Axiom™ Equine 670K SNP genotype data from 2,768 equids representing 20 horse breeds and one donkey breed. While assessing genome-wide runs of homozygosity (ROH) patterns, we observed an increased number of heterozygous calls in 167 purebred horses, which exhibited fewer ROH segments compared to F1 crosses, with 24 of them completely lacking any ROH segments. To further investigate this, we conducted a 4-fold genotype concordance analysis of replicate pairs on the same Axiom™ batch, between two different Axiom™ batches, between Illumina EquineSNP50 BeadChip® and between Illumina paired-end HiSeq 2000 whole genome sequencing data. Additionally, we used SNPolisher™ classification on data from the Axiom™ Equine 670K SNP array to evaluate the genotype performance of the 670,806 genome-wide SNPs. When comparing the overlapping SNPs between the different genotyping platforms, replicated pairs within the same Axiom™ batch showed the highest average genotype concordance (98.81%), followed by Illumina 50K (97.88%) and whole genome sequencing (96.84%). In contrast, re-genotyped horses with few ROH segments (i.e., replicates on different batches) showed the lowest concordance (93.52%). A lower pass rate was observed in one batch, suggesting a processing performance issue that contributed to the reduced concordance between batches. According to SNPolisher™ classification a total of 120,838 genome-wide SNPs were not recommended for reproducibility. After calling genotypes of the two different batches together in accordance with Axiom™ Best Practice (e.g. removing failing samples before the final genotyping) and excluding non-recommended SNPs, genotype concordance improved in all comparisons: same Axiom™ batch (99.84%), Illumina 50K (98.33%), whole genome sequencing (97.81%), and different Axiom™ batches (98.59%). Based on these findings, we recommend excluding horses exhibiting an unusually high number of heterozygous calls, using only SNPs with validated genotype performance, and accounting for batch effects when analyzing Axiom™ Equine 670K SNP genotype data from different batches.

**Article Summary:** In a European Equine diversity study including over 2,000 equids from 20 horse breeds and one donkey breed, we detected an unusually high number of heterozygous calls using the Axiom™ Equine 670K SNP array. Genotype concordance testing across different platforms showed that horses with fewer runs of homozygosity (ROH) had notably lower genotype concordance rates. By applying Axiom™ Best Practices and excluding SNPs with poor reproducibility, genotype concordance significantly improved. These findings highlight the critical need to validate SNP quality to ensure reliable genotypes for equine genomics projects and to account for batch effects when analyzing SNP data from different Axiom™ batches.

## Introduction

Single nucleotide polymorphism (SNP) arrays are a powerful genomic technology to simultaneously genotype thousands to hundreds of thousands genome-wide SNP markers [1]. This data resource has become essential in livestock genetics, with key applications including genomic selection [2], parentage testing [3], disease resistance [4], and genetic diversity studies [5]. By enabling the estimation of genomic estimated breeding values (GEBVs), SNP arrays improve selection accuracy and accelerate genetic gains for economically important traits such as milk yield, growth rate, and fertility [6, 7]. They also support genome-wide association studies (GWAS) and quantitative trait loci (QTL) mapping to identify genes associated with a desirable trait [8]. Additionally, SNP data ensures reliable pedigree verification and can be used to ascertain the genetic diversity within and between breeds, hereby supporting the management decisions to preserve endangered and native breeds [9]. To date, Illumina® (www.illumina.com) and Affymetrix™ (www.thermofisher.com) offer a wide range of customizable and species-specific arrays including cattle [10, 11], pig [12, 13], sheep [14, 15], goat [16], chicken [17], horse [18, 19] and honeybees [20]. Hence, these two genotype platforms can be considered as key players providing accurate, cost-effective and high-throughput genotyping services.

SNP array data is now widely used to estimate genomic inbreeding within breeds by identifying genome-wide patterns of homozygosity (ROH), which represent haplotypes that are identical-by-descent (IBD) [21]. The genomic inbreeding coefficient (F_ROH_) for an animal is calculated by dividing the sum homozygous segments (S_ROH_) by the total length of the genome. In livestock, previous studies have reported a high concordance between F_ROH_ and pedigree-based inbreeding coefficients (F_PED_) [22]. However, in our recent genetic diversity study of Franches-Montagnes (FM) horses, we observed notably lower F_ROH_ values compared to the corresponding F_PED_ values [23]. To further investigate this discrepancy, we re-analysed the Axiom™ Equine 670K SNP genotype data from a European equine diversity study, which included 2,768 equids across 21 breeds. Furthermore, we performed a 4-fold genotype concordance analysis using replicate sample pairs: (1) within the same Axiom™ batch, (2) between two different Axiom™ batches, (3) between Axiom™ and Illumina EquineSNP50 BeadChip® and (4) between Axiom™ and Illumina paired-end HiSeq 2000 whole genome sequencing data.

## Material and Methods

The Axiom™ Equine 670K SNP genotype dataset included 2,768 equids, representing 20 horse breeds and one donkey breed, and encompassed both previously published and unpublished data (Supplementary Table 1). Therefore, detailed information on most of the breeds can be found in the respective original publications. Unlike previous studies, the chromosomal SNP positions were determined based on EquCab3.0 reference genome [24]. For the ROH analysis, we excluded SNPs located on sex chromosomes or those without known chromosomal positions, resulting in a final dataset of 602,131 autosomal SNPs.

**Table 1.** Summary of Runs of homozygosity (ROH) analysis across 20 horse breeds and one donkey breed (*italicized*). The table includes the following information: average number of ROH (Ø N_ROH_), average length of ROH segments (Ø S_ROH_, in Mb), average length of ROH (Ø N_ROH_ in Mb), average genomic inbreeding (Ø F_ROH_), number of individuals with zero N_ROH_ and number of purebred individuals with less N_ROH_ compared to F1 outcrosses.

| Breed | N | $\bar{N}_{ROH}$ | $\bar{S}_{ROH}$ | $\bar{L}_{ROH}$ | $\bar{F}_{ROH}$ | $N_{ROH} = 0$ | $0 \leq N_{ROH} \leq 85$ |
| --- | --- | --- | --- | --- | --- | --- | --- |
| AKT | 51 | 176.06 | 231.75 | 1.22 | 10.16% | 1 | 6 ($\bar{N}_{ROH} = 54.33$) |
| AR | 184 | 254.49 | 349.89 | 1.37 | 15.34% | 0 | 0 |
| CT | 96 | 122.86 | 114.84 | 0.87 | 5.03% | 0 | 26 ($\bar{N}_{ROH} = 49.92$) |
| ES | 54 | 180.41 | 234.84 | 1.27 | 10.30% | 0 | 2 ($\bar{N}_{ROH} = 39.00$) |
| EXP | 278 | 291.23 | 420.52 | 1.37 | 18.44% | 10 | 10 ($\bar{N}_{ROH} = 23.50$) |
| FM | 853 | 171.48 | 279.66 | 1.63 | 12.26% | 1 | 4 ($\bar{N}_{ROH} = 64.25$) |
| FT | 156 | 196.28 | 346.45 | 1.76 | 15.20% | 0 | 1 ($\bar{N}_{ROH} = 49.00$) |
| HAF | 150 | 156.41 | 190.47 | 1.10 | 8.35% | 3 | 23 ($\bar{N}_{ROH} = 25.22$) |
| LIP | 335 | 196.29 | 280.17 | 1.38 | 12.28% | 1 | 16 ($\bar{N}_{ROH} = 50.69$) |
| LUS | 55 | 193.53 | 247.17 | 1.17 | 10.84% | 0 | 7 ($\bar{N}_{ROH} = 17.00$) |
| NOR | 207 | 136.76 | 158.96 | 1.03 | 6.97% | 5 | 42 ($\bar{N}_{ROH} = 35.24$) |
| POS | 30 | 146.67 | 184.68 | 1.20 | 8.10% | 1 | 0 |
| PRE | 15 | 196.07 | 362.38 | 1.79 | 15.89% | 0 | 0 |
| SA | 33 | 235.27 | 277.06 | 1.15 | 12.15% | 0 | 0 |
| SDH | 6 | 106.83 | 118.60 | 0.93 | 5.20% | 1 | 0 |
| SF | 297 | 193.82 | 282.74 | 1.44 | 12.40% | 1 | 5 ($\bar{N}_{ROH} = 59.40$) |
| SP | 4 | 191.25 | 186.67 | 0.94 | 8.18% | 0 | 0 |
| TB | 64 | 343.06 | 436.14 | 1.26 | 19.12% | 0 | 1 ($\bar{N}_{ROH} = 64.00$) |
| WB | 153 | 179.76 | 244.83 | 1.36 | 10.73% | 0 | 0 |
| ZH | 8 | 204.75 | 283.72 | 1.37 | 12.44% | 0 | 0 |
| *AHD* | 9 | 3.56 | 2.06 | 0.57 | 0.09% | 0 | 9 ($\emptyset N_{ROH} = 3.56$) |
| **Total** |  |  |  |  |  | **24** | **152** |

Runs of homozygosity segments were determined with an overlapping window approach implemented in PLINK v1.9 [25]. The following settings were applied: a minimum SNP density of one SNP per 50 kb; a maximum gap length of 100 kb; and a minimum length of homozygous segments of 500 kb (including more than 80 homozygous SNPs), while one heterozygote and one missing SNP was permitted per segment. The total number of ROH (N_ROH_), the total length of ROH segments (S_ROH_) and the average length of (L_ROH_) as well as F_ROH_, which was calculated by dividing S_ROH_ by the length of the autosomal genome (*L*_AUTO_ = 2280.92 Mb), were summarized for each breed.

Based on the ROH results of the horses, we re-genotyped five purebred FM and eight Lusitano (LUS) individuals, as these horses exhibited fewer ROH segments compared to F1 outcrosses. Additionally, two FM control horses with an average number of ROH segments were selected for re-genotyping. The genotyping and re-genotyping of the horses was outsourced to Neogen (www.neogen.com) using the same DNA sample. Quality of genotyping was considered

acceptable with a dish QC (DQC) ≥ 0.82 and QC call rate (CR) ≥ 97 according to Axiom™ best practice [26]. To further assess genotype concordance within the same Axiom™ batch, we also included 12 replicates (eight FM and three Warmblood horses). Furthermore, we used SNPolisher™ classification to evaluate the genotype performance of the 670,806 genome-wide SNPs across both Axiom™ batches including 27 replicates. Finally, we assessed genotype concordance between the Axiom™ batch and the Illumina EquineSNP50 BeadChip®, as well as between the Illumina paired-end HiSeq 2000 whole-genome sequencing data, including 25 and six FM horses, respectively. Therefore, we updated the chromosomal SNP position according to the EquCab3.0 reference genome and determined the number of overlapping SNPs between the Axiom™ Equine 670K SNP array and the two Illumina platforms containing 46,802 and 476,268 autosomal SNPs, respectively. Genotype concordance was calculated with Plink v1.9 [25] with merge-mode 6 command, which includes only non-missing calls present in both datasets.

## Results and Discussion

In our European diversity dataset, we observed 24 horses across nine breeds entirely lacking ROH segments. Additionally, 152 purebred horses originating from 12 different breeds exhibited fewer ROH segments (N_ROH_ < 85) compared to F1 outcrosses (Table 1). Consequently, only six horse breeds did not include any individuals with an increased number of heterozygous calls. In contrast, all genotyped donkeys displayed a high level of heterozygosity, as evidenced by the lowest number of ROH segments within the dataset. This unexpected high heterozygosity in donkeys may reflect a SNP ascertainment bias when calling donkey genotypes alongside horses. However, such biases have been previously linked with an increased number of homozygous calls for breeds that were underrepresented or not included in the SNP discovery panel [27]. The performance of the donkey samples may be improved by processing at least 96 samples separately, as recommended in the Axiom™ Best Practices guidelines. Moreover, it could be noticed that HAF horses associated with a low number of ROH segments also did not carry the respective homozygous allele state for the breed-specific chestnut coat color [28].

In Figure 1 (panels A-D, grey bars), we present the individual genotype concordance rates across the four comparison scenarios. The highest average concordance was observed within replicated pairs processed on the same Axiom™ batch (98.81%), followed by Illumina 50K array (97.88%), whole-genome sequencing (96.84%), and comparison across different Axiom™ batches (93.52%). Two replicated pairs within the same Axiom™ batch showed lower N_ROH_ values than F1 crosses (Figure 1, panel A). However, based on the replicate genotype set, these horses carried 109 and 154 N_ROH_ segments, respectively and exhibited the lowest genotype concordance in that comparison. Notably, the cross-batch comparison included re-genotyped horses with extreme low numbers of ROH segments (N_ROH_ ≤ 50), which were associated with reduced concordance rates (Figure 1, panel B). In contrast, horses with a moderate number of ROH segments (100 ≤ N_ROH_ ≤ 180) demonstrated higher genotype concordance. Additionally, we observed a lower pass rate in the batch containing horses with few ROH segments, suggesting a processing performance issue that contributed to the reduced concordance rates. In the Illumina 50K and whole genome sequencing (WGS) comparison we noticed one horse exhibiting 104 N_ROH_ segments, consistently demonstrating the lowest genotype concordance in both comparisons. This observation aligns with our previous findings, where this horse also showed low concordance when comparing F_ROH_ against F_PED_ [23].

**Figure 1.**
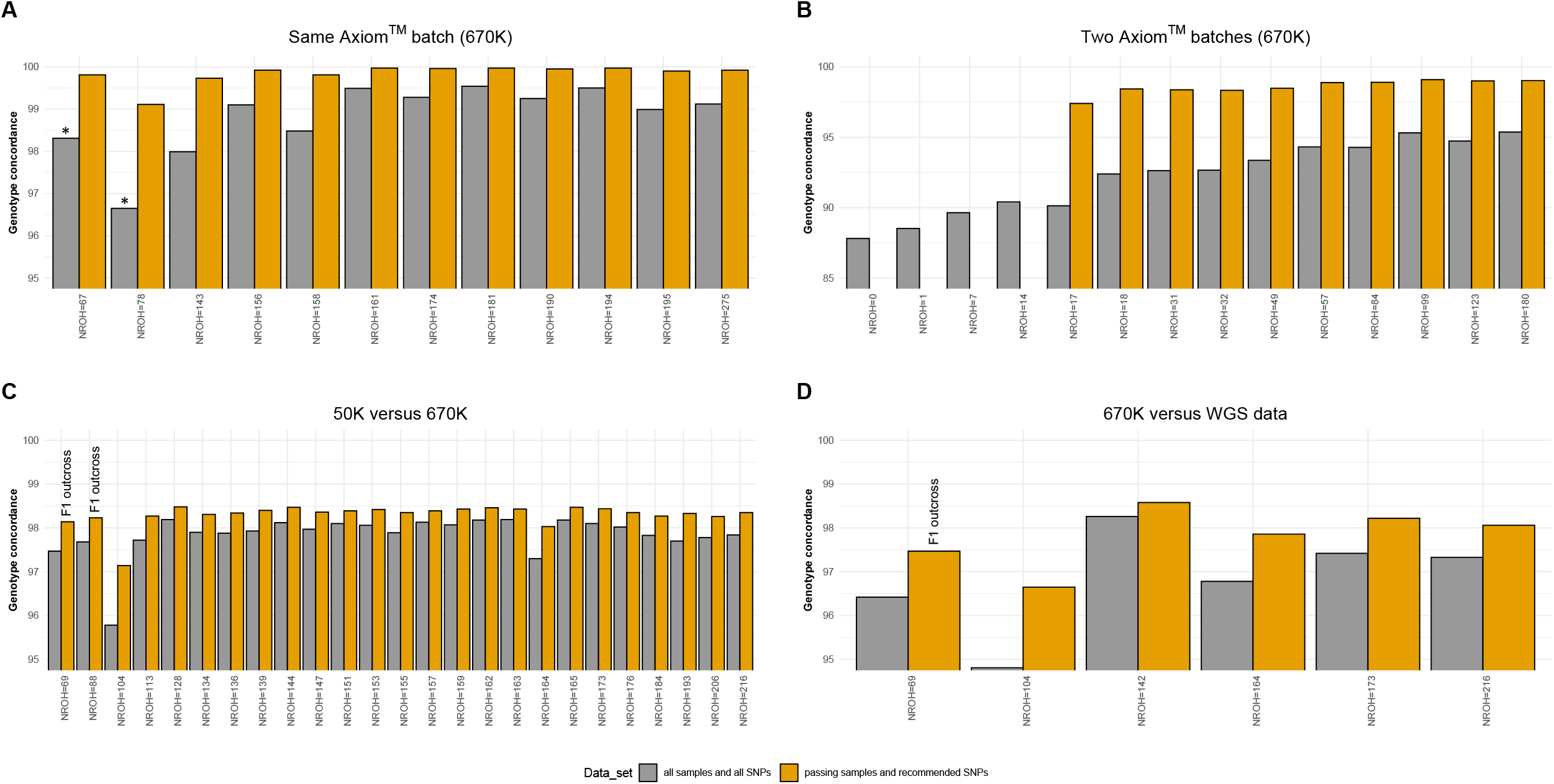
Four-fold genotype concordance analysis. Panels A-D illustrate genotype concordance across for scenarios - same Axiom™ batch, two Axiom™ batches, 50K versus 670K, 670K versus whole genome sequencing (WGS) - using two different sample and SNP sets. Each bar represents an individual horse, arranged from left to right by an increasing number of homozygosity segments (N_ROH_). The bar color indicates which sample and SNP set were used for genotype concordance analysis. Replicate pairs within the same Axiom™ batch exhibiting less N_ROH_ than F1 outcrosses are marked with (*), and F1 outcrosses are also highlighted.

During quality control (QC) of the two merged Axiom™ batches (192 individuals), we identified four additional failing samples including a re-genotyped FM horse exhibiting no ROH segments and three LUS horses carrying less than 14 ROH segments, which were therefore excluded from downstream analysis. Following this, we applied the SNPolisher™ classification to the genome-wide SNP data. The SNPs were allocated into six different categories (Supplementary Table 2). The vast majority (549,968 SNPs) fall into high quality categories (PolyHighResolution, NoMinorHom and MonoHighResolution), while a total of 120,838 SNPs were flagged as unreliable and not recommended. These unreliable SNPs were categorized as CallRateBelowThreshold (43,383 SNPs), OffTargetVariant (4,477 SNPs) and Other (72,978 SNPs) and were randomly distributed across the genome without any discernible pattern (Supplementary File 1). After calling genotypes of the two different batches together in accordance with Axiom™ Best Practice (e.g. removing failing samples before the final genotyping) and excluding non-recommended SNPs, genotype concordance improved in all comparisons (Figure 1, panels A-D, orange bars): same Axiom™ batch (99.84%), Illumina 50K (98.33%), whole genome sequencing (97.81%), and different Axiom™ batches (98.59%). However, concordance remained slightly lower than what was previously reported for Illumina 50K versus Illumina paired-end HiSeq 2000 data (98.50%) [29].

Excluding non-recommended SNPs prior to identifying ROH segments, most breeds displayed, on average, a similar number of N_ROH_ compared to the full SNP set (Figure 2A). Interestingly, CT, which included the second highest number of horses (26) with fewer ROH segments than F1 crosses, showed a significant increase in N_ROH_. In contrast, NOR, which had the highest number of horses (42) with N_ROH_ ranging from 1 to 85, showed no noticeable difference. Unexpectedly, EXP and TB, showing the highest average number of N_ROH_, exhibited fewer N_ROH_ when analyzing the edited SNP set. Furthermore, even after SNP editing, some breeds still included horses with N_ROH_ equal to zero, confirming previously observed QC results in FM and LUS horses, which clearly identified such horses as outliers. Some of these horses only showed a slight increase in N_ROH_, and on average, no significant difference was observed between the two SNP sets (Figure 2B). In contrast, for horses with N_ROH_ ranging from 1 to 85, a significant difference was noted between the two SNP sets. One TB horse stood out, exhibiting 64 N_ROH_ based on the full SNP set and 251 N_ROH_ using the edited SNP set, whereas, on average, TB horses had fewer N_ROH_ with the edited SNP set (Figure 2A).

**Figure 2.**
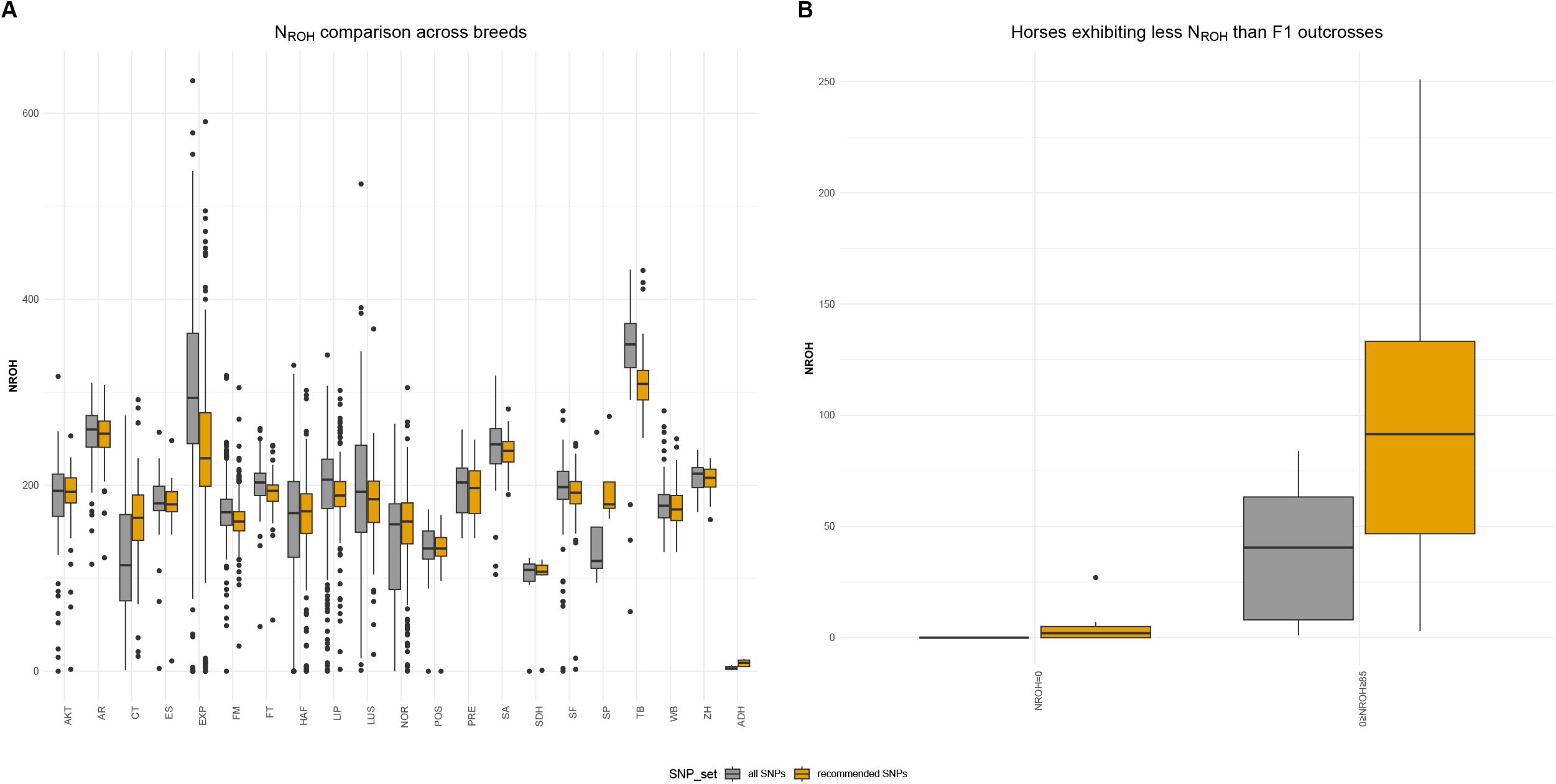
Runs of homozygosity (ROH) comparison between two SNP sets. (A) Number of ROH segments (N_ROH_) across 20 horse breeds and one donkey breeds, analyzed using two different SNP sets. (B) Comparison of SNP sets in horses exhibiting less N_ROH_ than F1 outcrosses including horses with zero N_ROH_.

## Conclusion

Our results demonstrate that the identification of ROH segments is a reliable indicator for assessing the genotype quality of individual horses, particularly when a low number of ROH segments is observed. Accordingly, we recommend excluding horses with an unusually high number of heterozygous calls and limiting downstream analyses to SNPs with validated genotype performance. However, if such low-quality individuals are present in the dataset, genotypes should be re-called after removing these horses to ensure high-quality genotype data. Additionally, we advise accounting for batch effects when analyzing 670K Axiom™ SNP data from different batches. Given the increased number of heterozygous calls observed in donkeys, we do not recommend calling donkey genotypes alongside horses.

## Data availability statement

Original and updated genotype information of all re-genotyped horses will be uploaded to Zenodo.

### Acknowledgements

We extend our sincere gratitude to our colleagues Thomas Druml (University of Veterinary Medicine Vienna), Vinzenz Gerber (University of Bern), Gabrielle Lindgren (Swedish University of Agricultural Sciences), Anne Ricard (INRAE), and Brandon Velie (University of Sydney) for generously sharing the Axiom™ Equine 670K SNP genotype data of their sampled breeds, which was essential to this study. Additionally, we appreciate the insightful discussions and collaborative spirit fostered during the 14^th^ International Havemeyer Foundation Horse Genome Workshop held in May 2024 at the University of Caen.

## Study Funding

This study was funded by the Swiss Federal Office for Agriculture (FOAG) under contract number 625000469 and Foundation Sur-La-Croix (internal contract number 6510263).

## Conflict of interests

Liz McInnis and Ali Pirani are employed by the manufacturer (Thermo Fisher Scientific) of Axiom™ Equine 670K SNP array, which may be perceived as a potential conflict of interest. However, the research was conducted objectively, and the authors declare that there are no commercial or financial relationships that could be construed as a potential conflict of interest.

